# Unlocking the Secrets of Amphibious Plant Adaptation: The Role of RNA Modifications

**DOI:** 10.1101/2024.11.25.625166

**Authors:** Mateusz Maździarz, Katarzyna Krawczyk, Joanna Szablińska-Piernik, Łukasz Paukszto, Monika Szczecińska, Paweł Sulima, Jakub Sawicki

**Affiliations:** University of Warmia and Mazury in Olsztyn, Department of Botany and Evolutionary Ecology

**Keywords:** epitranscriptome, N6-methyladenosine, 5-methylcytosine, pseudouridine, *Riccia fluitans*, flood stress

## Abstract

Post-transcriptional RNA modifications, such as N6-methyladenosine (m6A), 5-methylcytosine (m5C), and pseudouridine (Ψ), are critical regulators of plant development, stress responses, and environmental adaptation. The modifications m6A, m5C, and Ψ were insufficiently studied in plant stress responses. To investigate the epitranscriptomic landscape of these modifications, we employed direct RNA sequencing (DRS) to analyze native RNA from the amphibious plant *Riccia fluitans* grown in both aquatic and terrestrial environments. Our study revealed the presence of Ψ, m5C, and m6A modifications in *R. fluitans* transcriptomes from diverse environments. We observed correlations between the occurrence of these modifications and transcript length, as well as poly(A) tail length. Furthermore, we analyzed the expression of genes encoding Ψ synthases and methyltransferases to gain initial insights into the regulatory mechanisms underlying these processes in *R. fluitans*. By understanding how RNA modifications are regulated in response to environmental changes, we can unlock the secrets of the remarkable adaptability of amphibious plants. This knowledge could have eventually led to the understanding of the mechanisms of plant land colonization.

## Introduction

In the ever-evolving landscape of plant biology, the significance of post-transcriptional RNA modifications has emerged as a captivating area of research (Nachtergaele and He, 2016). These modifications, including N6-methyladenosine (m6A), 5-methylcytosine (m5C), and pseudouridine (Ψ), have been found to play pivotal roles in plant development, stress responses, and adaptation to diverse environmental challenges (Zhong *et al*., 2008; Cui *et al*., 2017; Anderson *et al*., 2018; Tang *et al*., 2020).

The m6A modification, which involves the methylation of adenosine at the nitrogen-6 position, is the most prevalent internal mRNA modification in eukaryotes (Linder *et al*., 2015). This modification is recognized by reader proteins, which mediate its downstream effects on RNA metabolism (Shen *et al*., 2019). In plants, m6A has been implicated in various aspects of RNA metabolism, including splicing, export, stability, and translation. Studies have revealed that m6A plays a crucial role in plant development, regulating processes such as flowering time, root development, and embryogenesis (Arribas-Hernández and Brodersen, 2020). The dynamic and reversible nature of m6A modification allows for rapid adjustments in gene expression, making it a versatile tool for plants to fine-tune their responses to changing environments. Therefore, m6A has emerged as a key player in plant stress responses, with evidence suggesting its involvement in responses to drought, heat, cold, salt, and pathogen stresses (Anderson *et al*., 2018; Liu *et al*., 2020; He *et al*., 2021; Yang *et al*., 2021; Hou *et al*., 2022). While m6A is primarily found in mRNA, other RNA modifications, such as m5C and Ψ, also seem play important roles in plant biology, but the studies on they role in stress responses and adaptation are limited (Liu *et al*., 2023; Sharma *et al*., 2023).

In contrast to m6A, which is predominantly found in mRNA, m5C is also present in other rRNA and tRNA (Sharma *et al*., 2023). This modification involves the methylation of cytosine at the carbon-5 position and is catalyzed by RNA methyltransferases (Tang *et al*., 2020). In plants, mRNA m5C modifications have been identified in various species, including *Arabidopsis thaliana* (David *et al*., 2017) and *Oryza sativa* (Tang *et al*., 2020). These modifications are involved in several essential processes, including plant development, stress responses, and long-distance mRNA transport (Guo *et al*., 2021). In *O. sativa*, the m5C methyltransferase OsNSUN2 mutants also exhibit shorter roots and increased sensitivity to heat stress. Notably, OsNSUN2 selectively methylates mRNAs involved in photosynthesis and detoxification, enhancing the plant’s heat tolerance (Tang *et al*., 2020).

Another RNA modification, Ψ is a ubiquitous RNA modification found in various RNA species, including messenger RNA (mRNA) in plants (Sun *et al*., 2019; Sharma *et al*., 2023). It is formed by the isomerization of uridine, resulting in a more rigid and stable RNA structure due to increased base stacking (Machnicka *et al*., 2013). In animals mRNA, Ψ has been shown to play crucial roles in regulating gene expression, mRNA processing, and stress responses by influence on mRNA splicing, stability, and translation, thereby affecting mRNA metabolism and gene expression (Liu *et al*., 2023), but the studies on plant mRNA are limited (Sun *et al*., 2019). Ψ modifications in mRNA have been implicated in various plant processes, including seed germination, root growth, and flowering time (Sun *et al*., 2019).

Understanding the roles of m6A, m5C, and Ψ modifications in plant adaptation and stress responses is crucial for developing strategies to enhance crop resilience and productivity in the face of climate change and other environmental challenges. However, these epitranscriptome modifications are poorly studied even in model plants, non-even mentioning non-model plants and their role in environmental adaptations.

The field of epitranscriptomics employs a variety of methods to detect RNA modifications like m6A, m5C and Ψ, each with its own set of advantages and drawbacks. For m6A, antibody-dependent methods such as MeRIP-seq, PA-m6A-seq, m6A-CLIP, and miCLIP are widely used (Yu *et al*., 2023). MeRIP-seq identifies m6A-enriched regions (Chen *et al*., 2019), while the others provide single-nucleotide resolution. Antibody-independent methods like MAZTER-seq and m6A- REF-seq rely on endoribonucleases (Garcia-Campos *et al*., 2019; Zhang *et al*., 2019), while chemical labeling methods like m6A-label-seq and m6A-SEAL offer alternative approaches (Shu *et al*., 2020; Wang *et al*., 2020). However, antibody bias, endoribonuclease motif preference, and labeling efficiency can limit their application. For m5C detection, bisulfite sequencing provides single-base resolution but suffers from low sensitivity for low-abundance modifications (Schaefer *et al*., 2009). Antibody-based methods similar to those used for m6A are also applied to m5C (Cui *et al*., 2017). Additionally, methyltransferase-dependent methods like Aza-IP and miCLIP enrich m5C-modified transcripts (Hussain *et al*., 2013; Khoddami and Cairns, 2013). Several techniques have been developed to detect Ψ within RNA molecules. Chemical modification with N-cyclohexyl-N′-(2- morpholinoethyl) carbodiimide metho-p-toluenesulfonate (CMCT) exploits the reactivity of Ψ, where CMCT modification blocks reverse transcription, allowing for identification of Ψ sites by comparing cDNA products generated with and without CMCT treatment (Carlile *et al*., 2014). Another method utilizes site-specific cleavage at Ψ residues using specific enzymes or chemicals, followed by radiolabeling of the cleaved fragments for identification (Meusburger *et al*., 2011). Mass spectrometry offers a direct detection method by analyzing the mass-to-charge ratio of RNA fragments, distinguishing Ψ from uridine based on their distinct masses (Yamauchi *et al*., 2016). Widely available SBS sequencing technologies coupled with specific chemical treatments or modifications allow for transcriptome-wide detection of Ψ (Marchand *et al*., 2020; Huber *et al*., 2023). However, challenges such as incomplete bisulfite conversion and overexpression of methyltransferases can lead to false positives. Furthermore, most of these methods require parallel controls, and the simultaneous detection of multiple modifications using NGS-based approaches remains a challenge.

Direct native RNA sequencing (DRS) is a cutting-edge method that allows sequencing of RNA molecules in their unaltered state, eliminating the need for reverse transcription into cDNA. This is achieved through Oxford Nanopore Technologies’ nanopore sequencers, which directly sequence native RNA strands as they traverse a protein nanopore (Nguyen *et al*., 2022; Wongsurawat *et al*., 2022). In contrast to traditional sequencing methods, DRS can identify RNA modifications, which are typically lost in widely used sequencing-by-synthesis (SBS) methods (Smith *et al*., 2019). This method has been employed to document nucleotide modifications and 3’ polyadenosine tails on RNA strands without additional chemical steps (Jain *et al*., 2022). DRS enables the analysis of native RNA strands without reverse transcription or amplification, thus avoiding biases introduced by these steps. Over recent years, the accuracy and throughput of direct RNA sequencing have significantly improved, reaching a point where it can offer valuable biological insights (Jain *et al*., 2022). For instance, it has revealed capping patterns in human mRNAs (Parker *et al*., 2020), detected novel Ψ sites in yeast (Marchand *et al*., 2020), and quantified changing modification levels under stress (Rudy *et al*., 2023). As the technology continues to advance, direct sequencing of full-length native RNA strands holds the potential to revolutionize transcriptomics (Yu *et al*., 2023).

While direct RNA sequencing has some limitations, it holds particular promise for studying non-model organisms exhibiting remarkable environmental adaptability, such as amphibious plants that adjust their morphology and physiology to thrive in fluctuating environments (Althoff *et al*., 2022; Koga *et al*., 2024). Comparative transcriptomics of amphibious plants grown submerged versus on land reveal differentially expressed genes involved in underwater acclimation like cuticle and stomatal development, cell elongation, and modified photosynthesis (Singh and Bowman, 2023; Maździarz *et al*., 2024). Moreover, comparative genomics between aquatic and terrestrial species identify genomic signatures enabling adaptation to submerged life, including changes in submergence tolerance, light sensing, and carbon assimilation genes (Shi *et al*., 2023). However, genomic resources for amphibious plants remain scarce, especially in the non-vascular evolutionary lineage. Understanding the RNA modifications in amphibious plants like *Riccia fluitans* can provide insights into their adaptive mechanisms to fluctuating environments. The aquatic liverwort *R. fluitans* is a close relative of model species *Marchantia polymorpha* (Bechteler *et al*., 2023), and serves as an excellent model for studying amphibious plants due to its remarkable adaptability (Althoff *et al*., 2022). *R. fluitans* exhibits exceptional environmental responsiveness, dynamically altering its morphology and physiology in response to its surroundings. When submerged, it develops thin thalli to maximize surface area for gas exchange. Upon emergence from water, it transforms into a land form with thicker thalli to reduce water loss and stockpiles starch in preparation for periodic drought (Althoff *et al*., 2022). This extreme plasticity enables *R. fluitans* to exploit both aquatic and terrestrial realms, demonstrating its ability to adapt to fluctuating conditions. Understanding the molecular mechanisms behind this adaptation in amphibious plants is an area of active research (Levin and Schuster, 2024; Maździarz *et al*., 2024).

The study aimed to investigate the impact of environmental changes on the epitranscriptome of the amphibious plant *R. fluitans*, focusing on three key RNA modifications: m6A, m5C, and Ψ. By employing cutting-edge DRS techniques, transcriptome profiling and bioinformatic analyses, the study sought to uncover the dynamic nature of the epitranscriptome and its role in the plant’s transition from terrestrial to aquatic environments. For this study we also adapt the ‘PRAISE’ method (Zhang *et al*., 2023*a*) to work with nanopore cDNA sequencing chemistry, enabling modified bases detection via bisulfite/sulfite treatment.

Identification of gene homologs involved in methylation (writers/erasers) and uridine modification (Ψ writers) in *R. fluitans* allowed us to verify the presence of a correlation between the expression of these genes and levels of RNA modification, bearing in mind that several studies drew conclusions about the epitranscriptome based solely on expression profiles.

## Materials and Methods

### Plant material, RNA extraction and RNA sequencing

Plant material (*R. fluitans* RF1 line) was obtained from an established axenic *in vitro* culture (Sawicki *et al*., 2023; Maździarz *et al*., 2024). Cultures were maintained in climate chambers at 24°C under long-day conditions (16-hour light/8-hour dark). After four weeks, one set of cultures was submerged in sterile distilled water, while another set remained unchanged. This experimental setup was replicated four times and continued for two weeks. Total RNA was extracted with the RNA Plant Mini Spin kit (Qiagen) following the manufacturer’s instructions. RNA quality and quantity were assessed using a Tapestation (Agilent) with the High Sensitivity RNA Screening Tape kit and a Qubit 4 fluorometer with the HS RNA Assay Kit.

Long-read native RNA libraries were prepared from 50 ng of poly(A)-tailed mRNA per sample using the Direct RNA Sequencing Kit SQK-RNA002 (Oxford Nanopore Technologies). Ribosomal RNA was removed with the NEBNext Poly(A) mRNA Magnetic Isolation Module (New England Biolabs). SuperScript III Reverse Transcriptase (Thermo Fisher Scientific) was used to synthesize the second strand of cDNA, forming RNA-cDNA hybrids. Sequencing adapters were then ligated using T4 DNA Ligase (New England Biolabs). Libraries were quantified with the Qubit dsDNA HS Assay Kit (ThermoFisher) and sequenced on a MinION MK1C device (ONT) with FLO-MIN 106 Flow Cells R.9.4.1 (ONT), prepared using the Flow Cell Priming Kit EXP-FLP002 (ONT). Raw reads were basecalled with Dorado 0.4.3 [https://github.com/nanoporetech/dorado] using the rna002_70bps_hac@v3 model on an NVIDIA RTX4090 GPU. The data is available in the ENA EMBL-EBI database under accession number PRJEB72691. Short-reads RNA sequencing libraries were constructed with the Truseq RNA library preparation kit (Illumina), incorporating the Ribo-Zero depletion step. Sequencing was performed on an Illumina NovaSeq 6000 platform at Macrogen Inc. (Seoul, Korea). Raw sequencing data has been deposited in the ENA EMBL-EBI database under accession number PRJEB72692.

### Expression profiling based on short-reads

RNA quality assessment was performed using FastQC software [https://www.bioinformatics.babraham.ac.uk/projects/fastqc/]. Following RNA-Seq, Illumina adapters and poly-A segments were excised with the Trimmomatic tool v.0.39 (Bolger *et al*., 2014). Reads shorter than 120 nucleotides (nt) and with an average quality score (PHRED) below 20 on leading and trailing sites were removed from the dataset. Next, high-quality reads were mapped to the draft genome (unpublished) using the STAR v.2.7.11a tool (Dobin *et al*., 2013). Obtained BAM files were used to create annotations with StringTie v.2.2.1 software (Shumate *et al*., 2022). StringTie aggregated individual GTF files from each sample and merged them into final annotations. Splicing variants of individual genes were obtained using the genomic annotations (GTF file). The count values for genes were then calculated by FeatureCounts v.2.0.6 with default parameters (Liao *et al*., 2014). A statistical test (based on a negative model) implemented in the DESeq2 v.1.42.0 (Love *et al*., 2014) R library was used to compare the expression profiles of water and land genes. An absolute logarithmic fold change (log2FC) greater than 0.54 and an adjusted p-value (padj) less than 0.05 were set as cut-off values for significantly differentially expressed genes (DEGs).

### Preprocessing long-reads

The POD5 signals from the MinION sequencer were first converted to fast5 format using the pod5- file-format program v.0.3.6 [https://github.com/nanoporetech/pod5-file-format/blob/master/docs/README.md]. Subsequently, basecalling was performed on the converted data using Guppy v.6.0.0 [https://nanoporetech.com/platform/technology/basecalling]. The quality of the received FASTQ files was evaluated using FastQC software, and these files were subsequently utilized for further analyses.

### m5C detection

Information on m5C methylation was obtained using the CHEUI software (Acera Mateos *et al*., 2024). Previously generated FASTQ files were mapped to the transcriptome sequence using the minimap2 software v.2.26 (Li, 2018) with the -ax map-ont parameter. The resulting bam files were then sorted using samtools v.1.7.2 (Danecek *et al*., 2021). The signal data was rescaled to the aligned sequences using the Nanopolish v.0.14.1 program [https://github.com/jts/nanopolish]. Preprocessing procedures assume two models: the first predicts modification within individual reads, second anticipates the stoichiometry and modification probability within transcriptomic sites. The RNA modifications that varied between terrestrial and aquatic environments were calculated using the CHEUI pipeline [https://github.com/comprna/CHEUI]. m5C modifications were classified as statistically significant when the probability exceeded 0.6. Sites present in both aquatic and terrestrial environments underwent differential analysis, where only those with pval_U < 0.05 and abs(stoichiometry diff) > 0.1 were deemed statistically significant.

### m6A detection

The detection, significance, and differential analysis of m6A modifications were conducted using CHEUI software with the same parameters as those used in the m5C section. The CHEUI per-site detection results were compared to NanoSPA (Huang *et al*., 2024) and m6Anet (Hendra *et al*., 2022). To compare the results between these programs, it was established that all detected site adenine modifications with a probability exceeding 0.6 would be analyzed. All-context (AC) methylation was detected by the CHEUI package, while the NanoSPA and m6Anet packages were identified as DRACH motifs. DRACH were identified as being more specific, referring to motifs located in specific sequence contexts. Conversely, AC motifs were identified as encompassing all possible motifs.

### Ψ **detection**

The transcriptome of *R. fluitans* was created using StringTie and gffread v.0.12.7 (Pertea and Pertea, 2020). Information on Ψ was extracted from previously generated FASTQ files and transcriptome using the nanopsU program v.1.0.0 (Huang *et al*., 2021*b*). Next, gaps created during the read merging process were removed, and features from all U sites were extracted using scripts from the nanopsU program. Only U sites with more than 20 reads were processed for further analysis. A Ψ region was considered reliable when the probability exceeded 0.8.

### Validation of m5C and Ψ

Ψ assessment and validation in mRNA were conducted using separate RNA extracts obtained from a different batch of plant material that had been submerged in sterile distilled water. The analysis was conducted using a modified sulfite/bisulfite treatment method - PRAISE method (Zhang *et al*., 2023*a*). Briefly, 6 µl total RNA (550 ng) was dissolved in 50 µl of a mixture of freshly prepared potassium sulfite and sodium bisulfite (50:50, molar proportion) with the addition of 100 mM hydroquinone which is a 100:1 final mixture of bisulfite/sulfite solution and hydroquinone. After 5 h of incubation at 70°C, the mixture was desalted by passing it through Micro Biospin 6 chromatographic columns twice (Bio-Rad, 7326200). The desulfonation step was carried out by incubation of desalted RNA with an equal volume of 1DM Tris–HCl (pH 9.0) at 75 °C for 30Dmin. The reaction was then immediately stopped by chilling on ice and followed by RNA precipitation with 3 volumes of ethanol (99.9%). The quantity of extracted RNA was measured with the Qubit^TM^ RNA HS Assay Kit. Next, ca. 400 ng of total RNA was reverse transcribed into cDNA and amplified to prepare the cDNA library following the protocol specified by Oxford Nanopore Technologies (ONT) for cDNA-PCR Sequencing Kit (SQK-PCS114). Adapter-ligated cDNA sequences were sequenced on the MinION flow cell (vR10.4.1) with MinION Mk1C sequencing device.

### Differential adenylation

The nanopolish v.0.14.1 [https://github.com/jts/nanopolish] was used to obtain information about the lengths of tails in individual transcripts. The quality of nanopolish output files was analyzed using nanotail v.0.1.0 [https://github.com/smaegol/nanotail]. Only observations with qc_tag "PASS" and "SUFFCLIP" were qualified for further analysis. Tails longer than 10 nucleotides were qualified for further analysis. Modification sites with a probability greater than 0.6 were used for poly(A) tail analysis. The Wilcoxon test carried out in the R was used to determine the significance of changes in tail lengths between transcripts with identified RNA modifications and other non-modified transcripts of the same gene. A p-value < 0.05 was considered as significant.

### Transcript length, alternative splicing, G/C content

Transcript length, splicing event and G/C content were compared between transcripts with identified RNA modifications and their non-modified variants for the same genes. Transcript length analysis was performed based on genomic GTF annotations. For Ψ, transcripts were categorized as modified if at least one site had a probability greater than 0.8. In contrast, for m5C and m6A, transcripts were classified as modified when at least one mutation had a probability exceeding 0.6. The significance of differences in transcript lengths was assessed using the Wilcoxon test. A p-value of < 0.05 was considered statistically significant. Alternative splicing events were generated from the previously obtained GTF file using the SUPPA v.2.4 program (Trincado *et al*., 2018). The number of events in transcripts with RNA modifications was then compared to the number of events in other transcripts of the same gene. Splicing events were categorized into the following groups: Skipping Exon (SE), Mutually Exclusive Exons (MX), Alternative 5’ Splice Site (A5), Alternative 3’ Splice Site (A3), Retained Intron (RI), Alternative First Exon (AF), and Alternative Last Exon (AL). GC content was calculated in the R script based on the FASTA file of the transcriptome. The Wilcoxon test was conducted once more, with a statistical difference defined as a p-value of less than 0.05. The proportions of modified nucleotides (Ψ, m6A, and m5C) to their respective unmodified bases (U, A, and C) in the transcript were calculated by dividing the number of detected modified nucleotides by the total number of the corresponding unmodified bases.

### Functional annotations

All DEGs, transcripts with significant modification profile changes were annotated using BlastP v.2.12.0 (McGinnis and Madden, 2004). Given the incomplete and uncharacterized status of many *M. polymorpha* gene symbols in databases, the identification of translated genes in *R. fluitans* was based on *A. thaliana* protein homology. A cut-off threshold of e-value < 10e-5 was set for blastp homology searching. The resulting gene signatures, including DEGs and other epitranscriptome candidates, were scanned for enrichment in Gene Ontology (GO) function annotations using the g:Profiler v.0.2.2 R library (Kolberg *et al*., 2023). Biological processes (BP), cellular components (CC), and molecular functions (MF) were assigned as ontological terms to essential genes. Enrichment analysis with a false discovery rate (FDR) cut-off of < 0.05 was employed to identify GO and pathway annotations regulated by differentially expressed genes.

### De-/methyltransferase and Ψ synthase family identification

The identification of two groups engaged in RNA modification started from a selection of *Arabodopsis* homologous sequences. The 44 and 20 genes encoded de-/methyltransferase and Ψ synthase identified in *A. thaliana* were blasted against the *R. fluitans* transcriptome. The Ψ synthase gene family encodes writers’ enzymes. The de-/methyltransferases were categorized based on their writer and eraser functions. Additionally, the best BLAST matches with e-value < e10-5 were used to search homologous with *Physcomitrella patens* and *M. polymorpha*. Translated coding sequences from four species were aligned using the MAFFT tool (Katoh and Standley, 2013)and organized into two separate datasets: de-/methyltransferases and Ψ synthases. Both alignments were used to build gene trees by the RAxML (Stamatakis, 2014) bootstrapping method with the GAMMA protein model.

### Visualization

The visualizations were created in the R environment v.4.3.1. The ggvenn v.0.1.10 library was used for Venn diagrams. ComplexHeatmap v.2.18.0 (Gu *et al*., 2016) was used to create heatmaps and upset plots. Visualization of the genetic trees was achieved through the use of the ggtree v.3.10.1 package (Yu, 2020). The rest of the visualizations were created using the ggplot2 v.3.5.1 package (Wickham, 2016).

## Results

### Unveiling Ψ in *R. fluitans*: land vs. water adaptations

Ψ modifications were detected in 311 and 824 transcripts in the land and water forms of *R. fluitans*, respectively. A total of 164 and 416 statistically significant Ψ sites were identified in the land and water forms of *R. fluitans*, respectively (Fig. 1C, Tabel S1,S2). Similarly, statistically significant changes were observed in 102 and 275 transcripts, respectively (Fig. 1D, Table S1,S2). Analyses were performed to assess the frequency of pseudouridylation in the transcripts. It was observed that some transcripts possess more potential pseudouridylation sites than others. The transcripts exhibiting the highest levels of pseudouridylation in the aquatic form of *R. fluitans* were CL.14105.3, coded Fatty acid/sphingolipid desaturase and unannotated mRNA - CL.24113.1, both containing eight potential pseudouridylation sites. (Table S1) In the terrestrial form, other unannotated transcript CL.3752.1 exhibited 19 potential Ψ sites, while CL.14105.3 showed 9 (Table S2). A functional analysis was subsequently performed to gain a deeper understanding of the biological processes in which pseudouridylated transcripts are involved. It was hypothesized that such information would be crucial to elucidating the role of this process in *R. fluitans*. In the land form, 193 significant processes were identified and classified into 48 biological processes (BP), 84 cellular components (CC), and 61 molecular functions (MF). In the water form of *R. fluitans*, 360 significant processes were detected, comprising 161 BP, 98 CC, and 101 MF. Among the statistically significant processes identified for both groups were photosynthesis (GO:0015979), response to stress (GO:0006950), and response to stimulus (GO:0050896) (Fig. 1E, Table S3,S4).

**Fig. 1.**
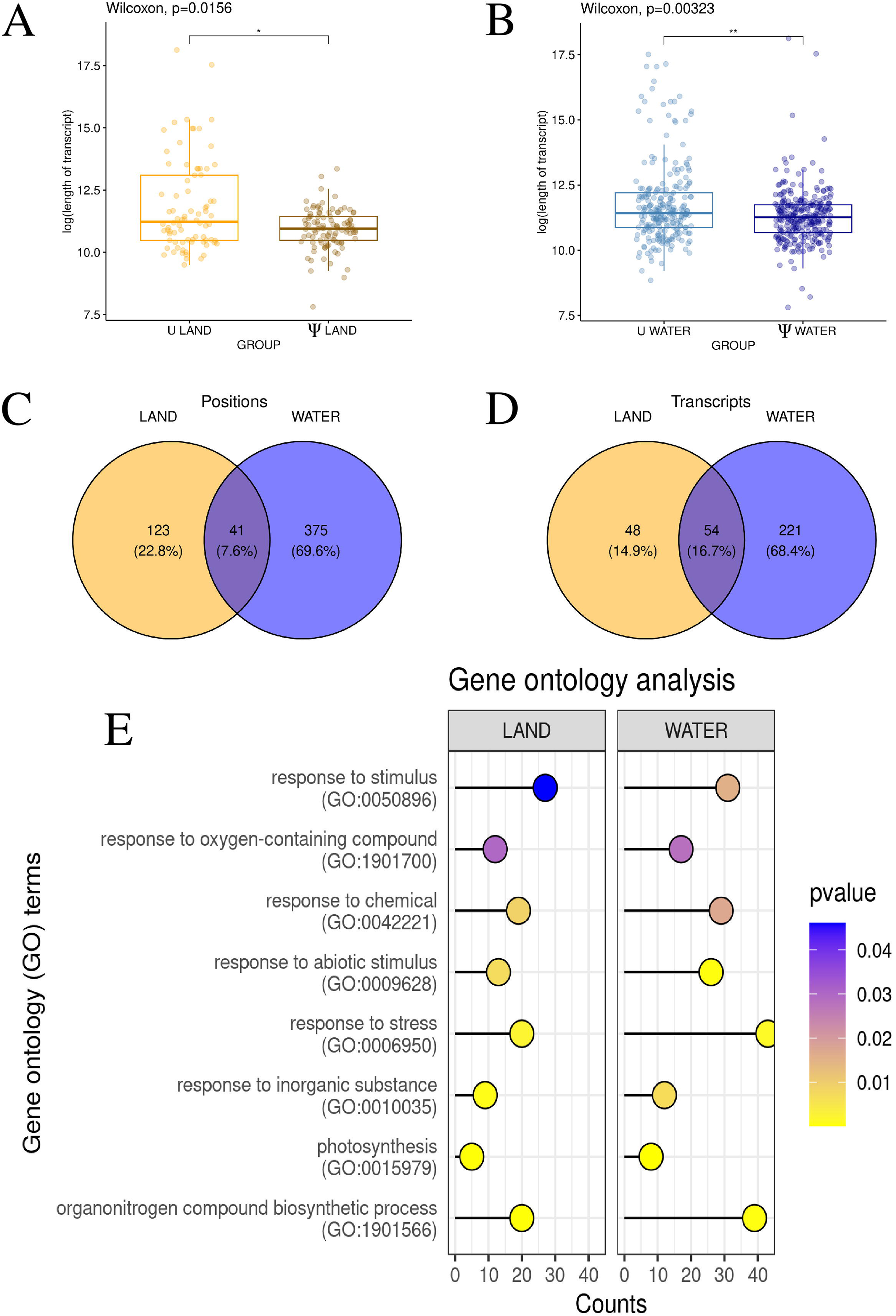
Ψ distribution and functional enrichment, with shorter transcript lengths in land and water *R. fluitans*. **A, B** – Boxplot charts depict the differences in log-transformed transcript length between Ψ-modified and non-Ψ-modified transcripts in two environments: land – A and water – B. **C, D** – Venn diagram illustrating the number of pseudouridylated sites – C, transcripts – D in land (yellow), water (blue), and both environments (dark blue). **E** – Lollipop chart illustrates the relationships between GO terms in the analysis, comparing land and water environments. The y-axis represents GO processes, while the x-axis shows the number of genes with detected pseudogenes involved in each process. Barplot colors are based on p-value significance.

The influence of Ψ on transcript length was subsequently investigated. Transcripts harbouring Ψ were observed to be shorter than transcripts of the same gene lacking a Ψ site. The difference in transcript lengths was found to be statistically significant for both the land (p = 0.0156) and the water forms (p = 0.00323) (Fig. 1A,1B, Table S5,S6). Since the analysis focused on introns and exons, it did not consider poly(A) tails, so we aimed to investigate how the length of this dynamic transcript structure changes. The length of the poly(A) tail and its correlation with transcripts containing Ψ were examined. The length of tails was found to be statistically significantly different between Ψ- containing transcripts and other transcripts of the same gene for both land (p = 0.003) and water (p < 0.001) (Fig. S1A,S1B, Table S7,S8). In both cases, a shortening of tails was observed in Ψ-containing transcripts. It was deemed reasonable to associate transcript length with the occurrence of alternative splicing. Alternative splicing, which can influence transcript length, was investigated. An analysis of pseudouridylated transcripts revealed the presence of Ψ sites near their splicing events, including 24 for alternative 3’ splicing (A3), 11 for alternative 5’ splicing (A5), 9 for alternative first exon (AF), 5 for alternative last exon (AL), 64 for retained introns (RI), and one for skipped exons (SE). In contrast, other transcripts of the same gene exhibited a distribution of splicing patterns with 29 (A3), 23 (A5), 28 (AF), 25 (AL), 9 (MX), 32 (RI), and 1 (SE) splicing events identified (Fig. S1C, Table S9).

The distribution of nucleotides in modified transcripts was also under consideration. It was assumed that the G/C content and the ratio of modified to unmodified nucleotides within the transcript would be sufficient to assess whether modified transcripts were more prone to modification due to a higher frequency of the nucleotide undergoing modification. A statistically significant difference in G/C content was found between Ψ-containing transcripts and other transcripts of the same gene in both terrestrial and aquatic forms (p-value < 0.001). The percentage of Ψ relative to all uridines in the transcript was determined to be 0.001% in the land form and 0.002% in the water form (Fig. S2A,S2B, Table S10,S11).

To confirm the reliability of the high throughput direct RNA sequencing results, the additional validation of RNA modification was performed. The localization of Ψ on transcripts was validated using the PRAISE method, which identified 42 positions in the aquatic form of *R. fluitans* where Ψ had previously been detected through direct RNA sequencing (Fig. 2)

**Fig. 2.**
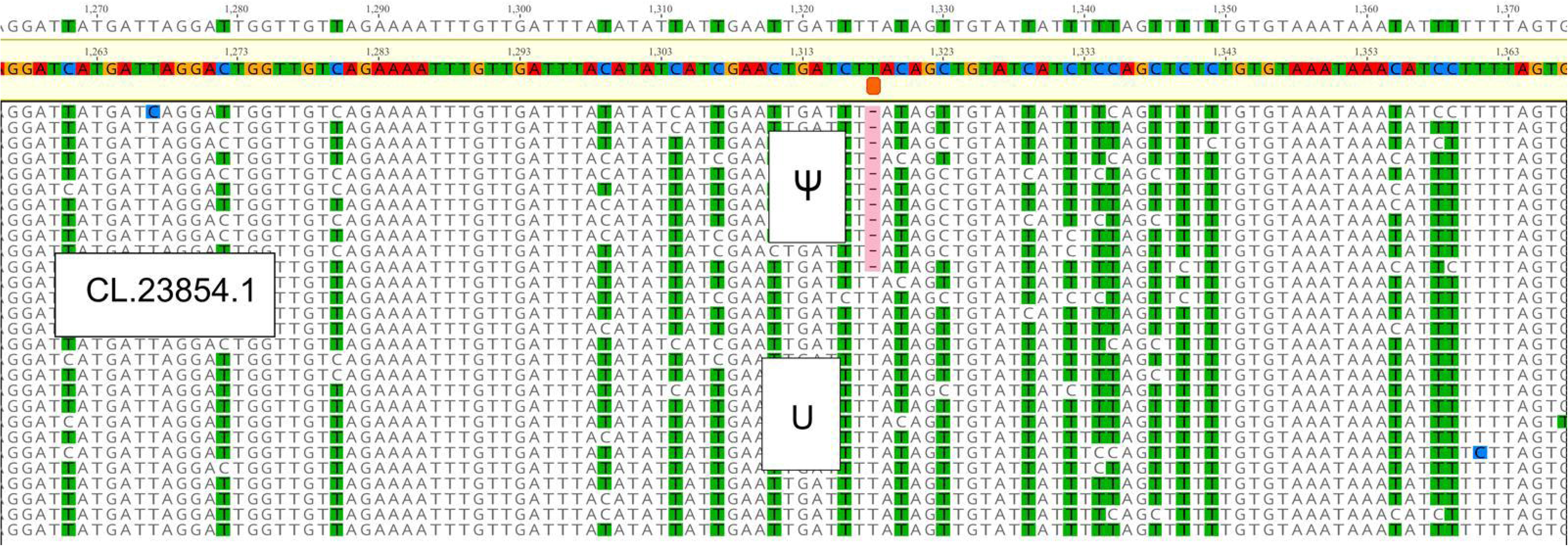
Visualization of CL.23854.1 transcript sequenced using PRAISE method adapted to nanopore sequencing technology. Presence of Ψ resulted in deletion at position 1318 which is congruent with NanoSPA prediction (probability 0.875).

### The impact of m5C methylation on transcript elongation in *R. fluitans*

AC analysis of m5C methylation revealed 26,330 putative methylation sites in *R. fluitans* habited in terrestrial environments and 85,295 growing in aquatic conditions. A total of 3,128 sites with a probability of greater than 0.6 were identified in land and 9,897 in water ecotypes (Fig. 3E,3F, Table S12,S13). To comprehensively analyze and compare the impact of the modification, similar analyses were conducted for m5C as for Ψ. Analysis of transcript length between transcripts with detected m5C and transcripts that had cytosine at this position revealed significantly different patterns in the aquatic environment (p = 0.009) (Fig. 3D, Table S14). Transcript elongation was observed in the aquatic environment with m5C, while there was no statistically significant difference in the terrestrial environment (p = 0.053) (Fig. 3C, Fig. S15). It was found that poly(A) tail lengths were shorter in transcripts with detected modification in water form (p < 0.001) (Fig. S3B, Table S16). However, no significant differences in the length of the poly(A) tail were detected in the terrestrial form (p = 0.45) (Fig. S3A, Tabel S17). Alternative splicing analysis was performed again to identify the type of event in transcripts burdened with modification. Splicing events were found to occur at a frequency of 260 (A3), 243 (A5), 56 (AF), 19 (AL), 5 (MX), 346 (RI), and 49 (SE) in transcripts with m5C. In contrast, a frequency of 83 (A3), 46 (A5), 40 (AF), 27 (AL), 7 (MX), 78 (RI), and 28 (SE) was observed in transcripts of the same gene without modification. (Fig. S3C, Table S18)

**Fig. 3.**
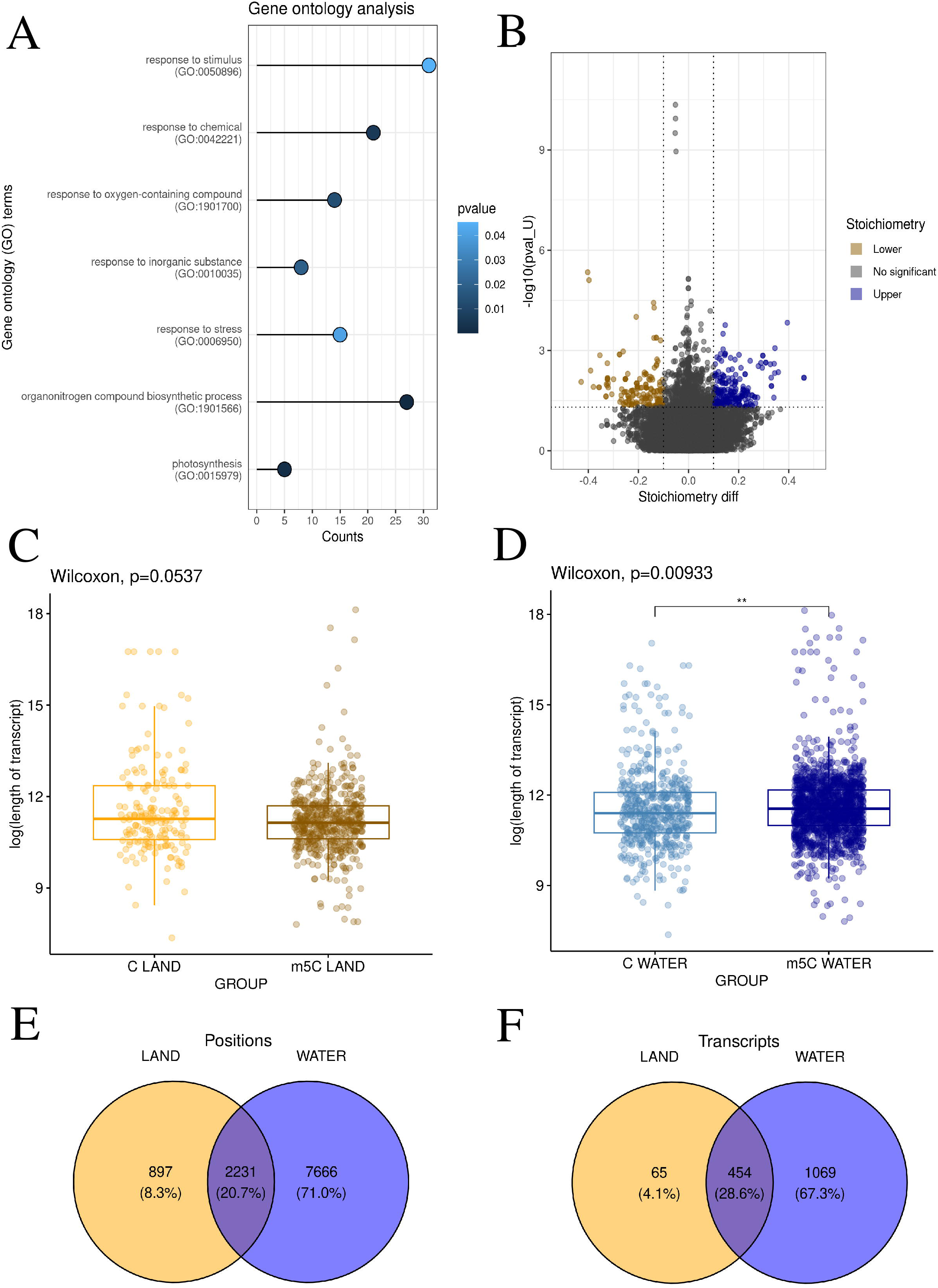
m5C methylation patterns and differential transcript elongation in land and water *R. fluitans*. **A** – Lollipop chart shows selected statistically significant GO processes on the y-axis, while the x-axis represents the number of m5C-modified genes involved in each process. p-values are represented by a blue scale, where dark blue indicates lower p-values. **B** – Volcano plot displays the methylation difference (stoichiometry diff) for each site. The x-axis represents the magnitude of these differences, while the y-axis shows the negative log-transformed pval_U. Sites with statistically significant methylation changes are highlighted by orange (lower) and blue (upper) color. The horizontal dashed line indicates the adjusted p-value cutoff (0.05). The vertical dashed lines represent a stoichiometry diff threshold (abs(0.1)). **C, D –** Boxplot charts depict the differences in log-transformed transcript length between m5C- modified and non-m5C-modified transcripts in two environments: land – A and water – B. **E, F –** Venn diagram illustrating the number of methylated sites (E) and transcripts (F) in land (yellow), water (blue), and both environments (dark blue).

To examine the nucleotide composition in transcripts, analyses of G/C content and the ratio of modified cytosines to unmodified cytosines within the transcriptome were conducted once more. Transcripts with detected m5C modification were found to be significantly different in G/C content from other transcripts of the same gene in both land (p<0.001) and water (p<0.001). Lower G/C content was observed in transcripts with detected modification. The proportion of m5C to total cytosines in the transcript was determined to be 0.08% under aquatic conditions and 0.03% under terrestrial conditions (Fig. S2C,S2D, Table S19,S20) .

Next, the CHEUI software was employed to perform a differential analysis between the positions detected in both groups. It was decided to utilize this option to observe in detail how specific positions change under the influence of stoichiometry. The 21 532 common sites were qualified for differential analysis. 394 sites were qualified as statistically significant, of which 164 were classified as lower and 230 as upper (Fig. 3B, Table S21). Statistically significant sites were involved in the gene ontology processes response to stimulus, response to stress, response to oxygen-containing compounds, and response to abiotic stimulus (Fig. 3A, Table S22).

### Analysis of m6A methylation motifs reveals distinct patterns in land and aquatic environments

A comprehensive analysis of m6A methylation across plants occupying different environments uncovered 33,175 land-based and 112,823 aquatic methylation sites. The analysis of AC motifs conducted with the CHEUI program revealed 16,681 statistically significant positions in the aquatic habitats of *R. fluitans*, while 4,730 significant positions were identified in terrestrial environments (Fig. 4E, Table S23,S24). The analysis revealed the presence of these positions across 540 transcripts in the terrestrial form and 1,648 transcripts in the aquatic form of *R. fluitans* (Fig. 4F). Transcript lengths were compared between transcripts harboring m6A modifications and those without m6A modifications from the same genes. No significant differences in transcript lengths were revealed when comparing transcripts harboring m6A versus non-m6A transcripts from the same genes (p = 0.15 for land, p = 0.06 for water) (Fig. 4C,4D, Table S25,S26). Tail lengths were not significantly changed in the terrestrial environment (p = 0.08) (Fig. S4A, Table S28). However, a shortening of poly(A) tail lengths was observed in transcripts with m6A modifications in the aquatic environment (p < 0.001) (Fig. S4B, Table S27). Splicing events were also observed in transcripts with m6A modification 284 (A3), 244 (A5), 58 (AF), 20 (AL), 6 (MX), 375 (RI), and 51 (SE) compared to other transcripts of the same gene 94 (A3), 78 (A5), 77 (AF), 35 (AL), 7 (MX), 90 (RI), and 40 (SE) (Fig. S4C, Table S29).

**Fig. 4.**
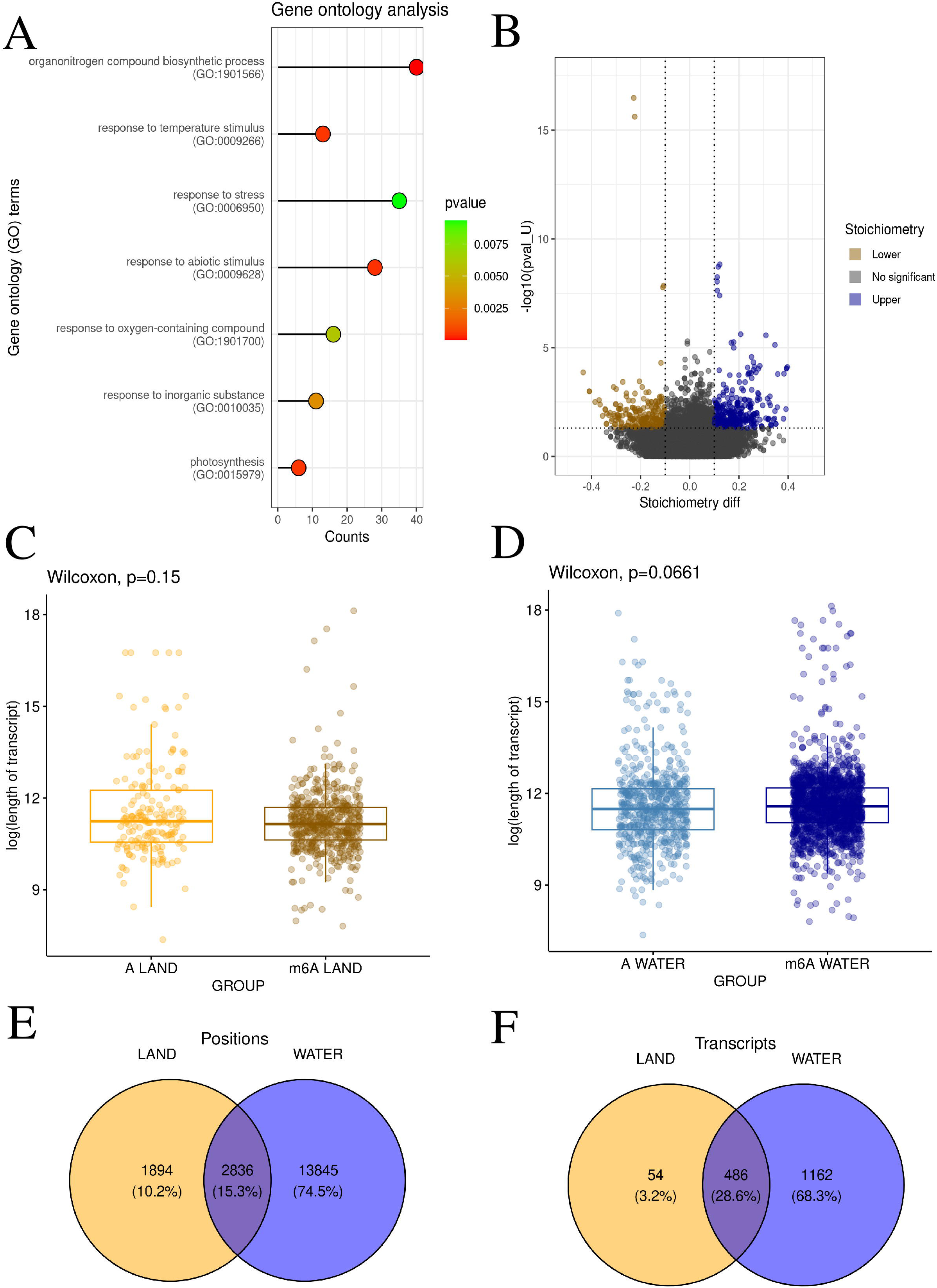
Distinct m6A methylation motifs and shared transcripts revealed in land and aquatic *R. fluitans*. **A –** Lollipop chart shows selected statistically significant GO processes on the y-axis, while the x-axis represents the number of m6A-modified genes involved in each process. The p-values are represented by a blue scale, where dark blue indicates lower p-values. **B** – Volcano plot displays the methylation difference (stoichiometry diff) for each site. The x-axis represents the magnitude of these differences, while the y-axis shows the negative log-transformed pval_U. Sites with statistically significant methylation changes are highlighted by orange (lower) and blue (upper) color. The horizontal dashed line indicates the adjusted p-value cutoff (0.05). The vertical dashed lines represent a stoichiometry diff threshold abs(0.1). **C, D –** Boxplot charts depict the differences in log-transformed transcript length between m6A- modified and non-m6A-modified transcripts in two environments: land – A and water – B. **E, F –** Venn diagram illustrating the number of methylated sites (E) and transcripts (F) in land (yellow), water (blue), and both environments (dark blue).

A comprehensive analysis of nucleotide distribution revealed that the GC content was downgraded for transcripts with m6A modifications in both terrestrial and aquatic environments (p < 0.001). The percentage of m6A relative to all adenines in the all transcript was determined to be 0.04% under terrestrial conditions and 0.1% under aquatic conditions (Fig. S2E,S2F, Table S30,S31). A differential methylation analysis targeting 26,981 sites identified in both groups revealed that 612 of these sites exhibited significant differences in m6A methylation. Of these, 247 m6A positions were found to display lower stoichiometry, while 365 sites indicated higher stoichiometry (Fig. 4B, Table S32). These significant sites were associated with 195 genes, which were found to be functionally enriched in processes related to "response to stimulus", "response to stress", "response to oxygen-containing compound", and "photosynthesis" (Fig. 4A, Table S33).

Motif enrichment analysis utilizing DRACH and AC motifs identified 4 and 39 transcripts, respectively, that were common between the terrestrial and aquatic forms of *R. fluitans*. Furthermore, 156 statistically significant m6A RNA modification sites were identified in the aquatic *R. fluitans* species by the NanoSPA program, and 173 such sites were identified by the m6Anet software. In the terrestrial *R. fluitans* ecotype, the 68 and 27 statistically significant m6A modification sites were identified by both programs, respectively (Fig. S5).

### Overlap of epitranscriptomic modifications in *R. fluitans*

A comprehensive analysis of shared epigenomic features was conducted using multiple software tools and methylation detection techniques. The results revealed that m5C and m6A modifications exhibited the highest degree of overlap, likely due to the predominance of the AC method. The DRACH motif was not the primary focus in the detection of modifications (Fig. S6A-F). In total, 304 transcripts were found to contain all three investigated modifications. Additionally, a smaller number of unique transcripts were identified between Ψ and m5C (3, 0.2%) and Ψ and m6A (7, 0.4%) (Fig. 5B).

**Fig. 5.**
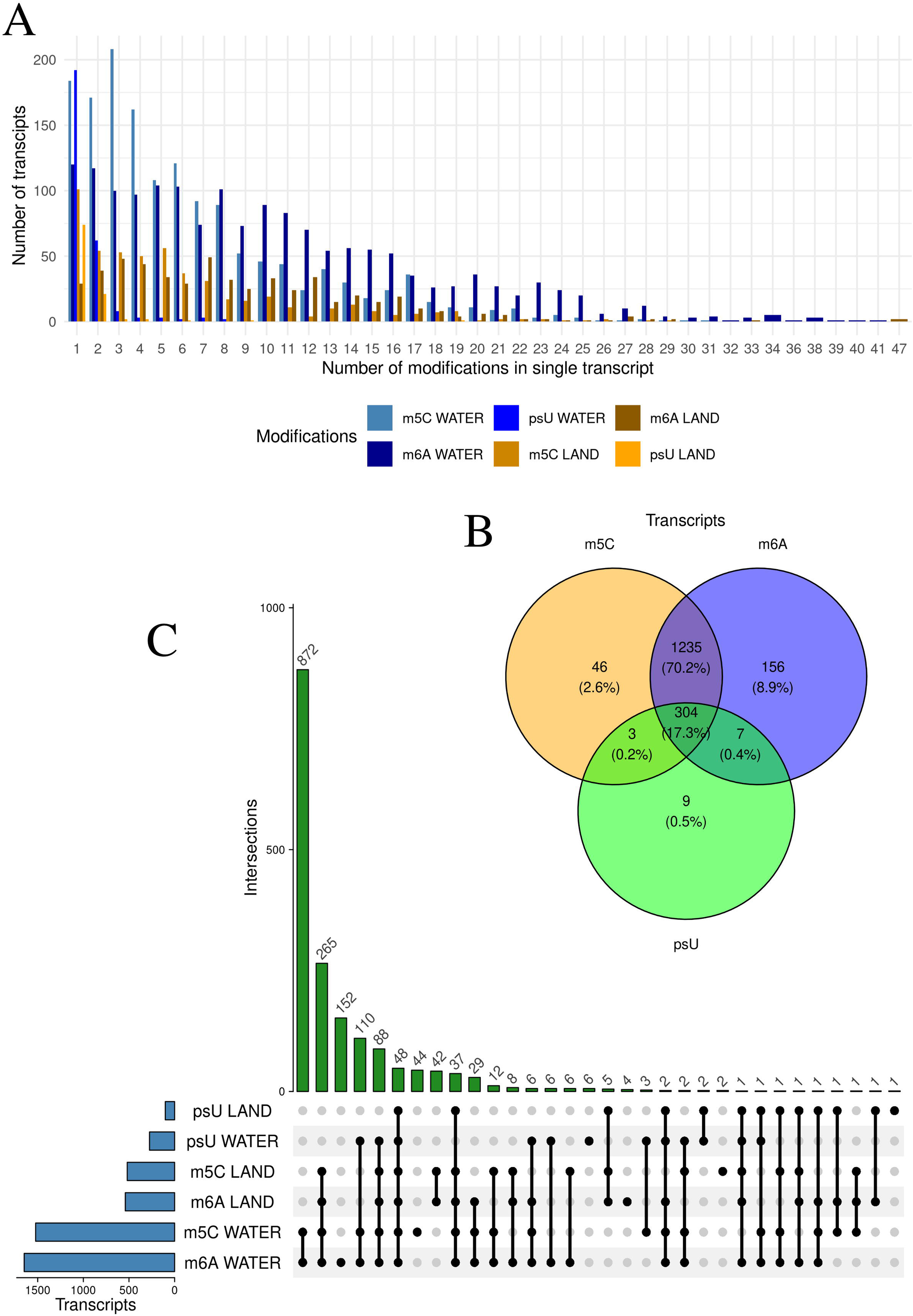
Transcript modification profile: distribution, overlap, and co-occurrence. **A** – The barplot illustrates the differences in distribution between transcripts. The y-axis represents the number of transcripts in which a specific number of modifications were detected, while the x-axis differentiates the number of modifications in a single transcript. **B –** The Venn diagram illustrates the relationships between the occurrence of modifications in transcripts for m5C (yellow), m6A (blue), and Ψ (green). **C –** The upset plot displays the transcripts shared across all modifications. The blue bar plots represent the number of transcripts for each modification, while the green ones indicate those that are jointly compatible, marked by filled black dots.

A summary of the modifications in the *R. fluitans* transcriptome indicated that a single position Ψ modification was the most prevalent, occurring in 192 aquatic transcripts and 74 terrestrial transcripts. For m5C, the highest abundance of transcripts (208) was observed with three modifications in the aquatic form of *Riccia*, whereas in the terrestrial form, the highest frequency of transcripts was associated with a single modification. For m6A, the aquatic form of *Riccia* exhibited the highest number of transcripts (120) with a single modification, while the terrestrial samples showed the highest number of transcripts (49) with up to seven modifications (Fig. 5A). The aquatic condition induced the highest number of m6A and m5C modifications within 872 transcripts expressed in *R. fluitans*. Other 265 unique transcripts were modified by methylation of cytosine and adenine in both environments. Finally, all three RNA modifications (Ψ, m6A, and m5C) were detected in 48 transcripts across both habitats of *Riccia* (Fig. 5C).

### Comparative analysis of de-/methyltransferase and Ψ synthase gene families in the liverwort *R. fluitans*

Beside the epitranscriptomic profiling, the research examined the expression of 20,840 *R. fluitans* genes using Illumina sequencing data. Of these, 1,717 exhibited statistically significant gene expression changes between isolates extracted from plants grown in terrestrial and aquatic habitats. Among the DEGs, 978 exhibited downregulation (higher expression in *Riccia* terrestrial form) and 739 were upregulated (higher expression in *Riccia* aquatic form) (Table S34). We aimed to determine whether differences in potentially pseudouridylated or methylated positions were attributable to variations in the expression of genes encoding proteins that function as RNA de-/methyltransferases and Ψ synthase complexes. The threshold for the significance of differences in expression was lowered (log2FC > 0.54). Nevertheless, no differences in the expression of molecules responsible for these processes were detected between the aquatic and terrestrial forms of *R. fluitans*. The correlation between expression in the aquatic and terrestrial environments for methyltransferases was R = 0.93 with a p-value < 0.05, indicating a very strong relationship (Fig. 6B). A similar pattern was observed for the correlation of Ψ synthases, with an R-value of 0.94 and a p-value < 0.05 (Fig. 7B).

**Fig. 6.**
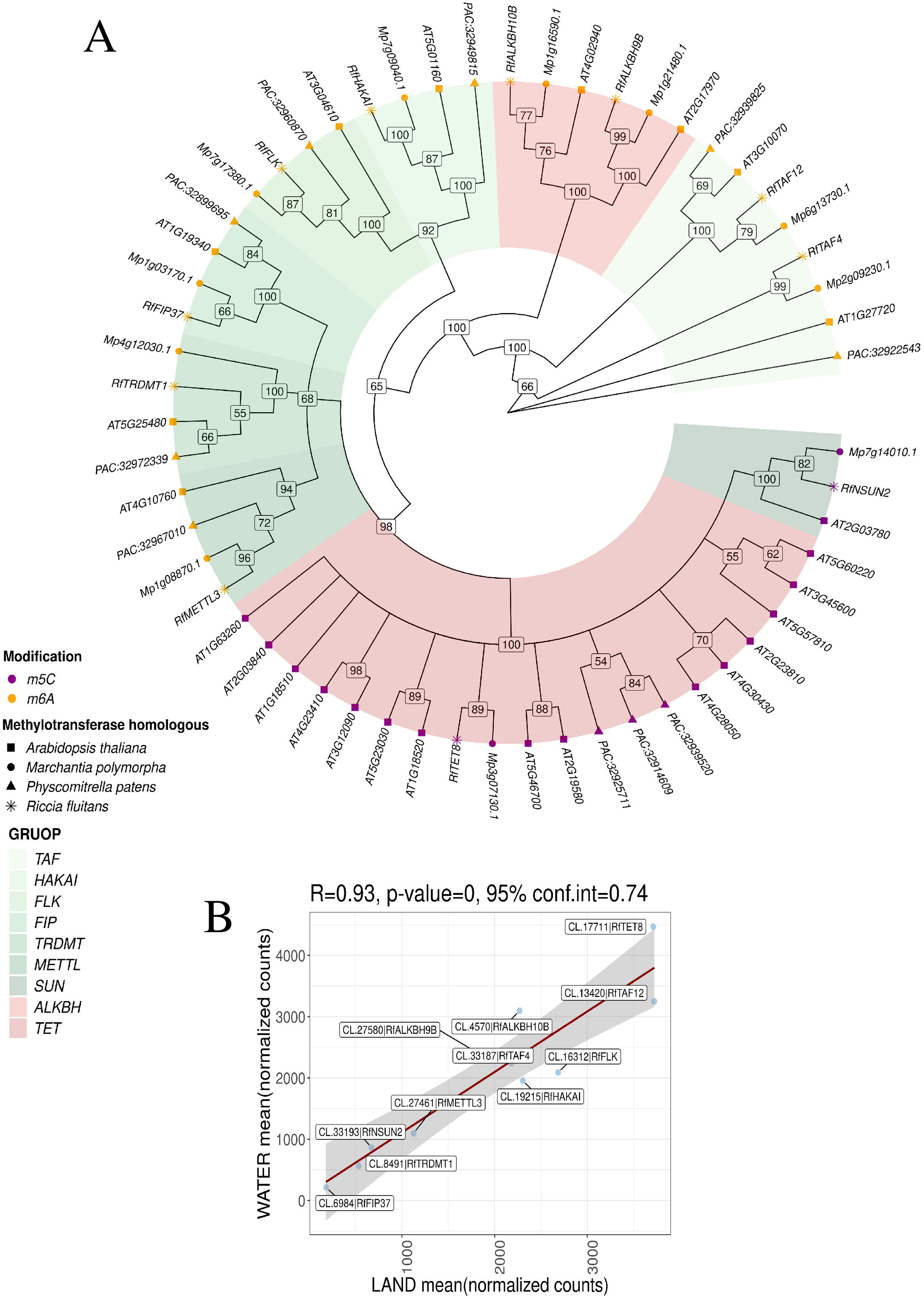
Evolutionary relationships of methyltransferase genes across *A. thaliana*, *M. polymorpha*, *P. patens*, and *R. fluitans* **A –** Phylogenetic tree of methyltransferase genes. The tree was constructed for four species: *A. thaliana* (square), *M. polymorpha* (circle), *P. patens* (triangle), and *R. fluitans* (star). Branch tips colored purple indicate genes involved in the m5C methylation process, while orange tips indicate those involved in the m6A methylation process. Green-colored clades suggest that the encoded proteins function as "writers" of these modifications, whereas red-colored clades suggest that they function as "erasers". **B** – The dotplot comparing the mean of normalized counts for identified methyltransferases in the aquatic form of *R. fluitans* (y-axis) and the terrestrial form of *R. fluitans* (x-axis). The red line highlights the Pearson correlation between the two forms. The R value, p-value, and confidence interval (conf.int) are displayed in the upper left corner. The gray bands around the line represent the standard error of the regression line.

**Fig. 7.**
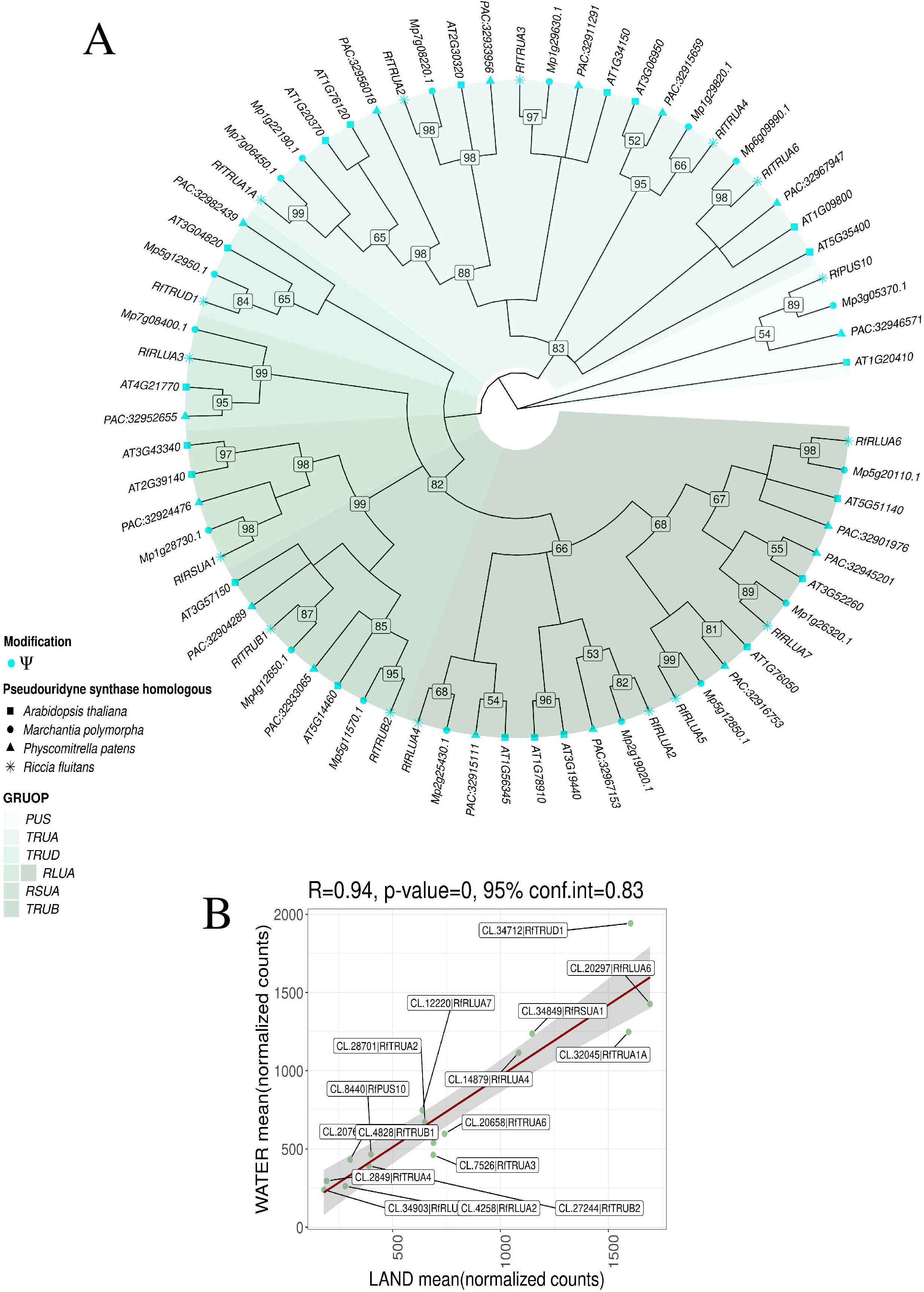
Evolutionary relationships of. Ψ synthases genes across *A. thaliana*, *M. polymorpha*, *P. patens*, and *R. fluitans* **A –** Phylogenetic tree of Ψ synthase genes. The tree was constructed for four species: *A. thaliana* (square), *M. polymorpha* (circle), *P. patens* (triangle), and *R. fluitans* (star). **B** – The dotplot comparing the mean of normalized counts for identified Ψ synthases in the aquatic form of *R. fluitans* (y-axis) and the terrestrial form of *R. fluitans* (x-axis). The red line highlights the Pearson correlation between the two forms. The R value, p-value, and confidence interval (conf.int) are displayed in the upper left corner. The gray bands around the line represent the standard error of the regression line.

Forty-four *A. thaliana* genes encoding de-/methyltransferases and twenty genes encoding Ψ synthases were used to identify homologous genes in *R. fluitans*. Blast searches identified eleven de-/methyltransferases and sixteen Ψ synthases unigenes. Among the methyltransferases that function as writers, a homology scan revealed 8 protein-coding genes expressed in the liverwort *R. fluitans* (*RfHAKAI*, *RfFLK*, *RfNSUN2*, *RfTRDMT1*, *RfTAF4*, *RfMETTL3*, *RfFIP37*, *RfTAF12*). Conversely, three genes (*RfALKBH10B*, *RfALKBH9B*, and *RfTET8*) were identified as essential for RNA demethylation, acting as erasers. In a comparative analysis of TET, *Arabidopsis* contains a substantial number of members, specifically 15 TET genes. In contrast, the liverwort *R. fluitans* exhibits only a single homologue of the Ten Eleven Translocation (TET) 8 gene. In the closely related species *M. polymorpha*, a unique homologue of the *TET8* gene has also been identified. Our analysis revealed that the moss *P. patens* possesses three distinct homologues within the TET gene family. Most gene clusters were represented by only one ortholog of *Arabidopsis*, *Marchantia*, *Physcomitrella*, and *Riccia*. Interestingly, the two erasers, *RfALKBH10B* and *RfALKBH9B*, along with the writer *RfNSUN2*, lacked homologous sequences in *P. patens*. Each de-/methyltransferase gene in *R. fluitans* (*Rf*) exhibited the closest relationship with homologous genes in *M. polymorpha* (*Mp*) (Fig. 6A, Table S35).

The research confirmed the occurrence of 16 genes encoded Ψ synthase enzymes in *R. fluitans*. All members of the six known subfamilies — RluA, RsuA, TruA, TruB, PUS10, and TruD — previously described in *A. thaliana* and *Zea mays*, were also identified in the transcriptome of *R. fluitans*. The most prominent subfamily, *RluA*, was divided into seven members, with the *RLUA3* gene distinctly separated from the other *RLU* genes. The moss (*Physcomitrella)* and liverworts (*Riccia* and *Marchantia*) each contained one representative within the *RfRLUA1/RfRLUA2* clade, whereas *A. thaliana* possessed both gene copies. Additionally, *A. thaliana* was found to possess two gene copies of At*RSUA1* and At*RSUA2*, whereas the other analyzed species displayed only a single representative. Similarly, within the *TRUA* clade, both *AtTRUA1A* and *AtTRUA1B* clustered with two copies from *Marchantia* and one *Riccia* homolog (*RfTRUA1A*) (Fig. 7A, Table S35).

## Discussion

The epitranscriptome of *R. fluitans* exhibits remarkable plasticity in response to changing environments, particularly the shift from terrestrial to aquatic habitats (Althoff *et al*., 2022; Maździarz *et al*., 2024). The frequency of RNA modifications, including m6A, m5C, and Ψ, increases substantially in the aquatic form, suggesting their crucial role in adaptive responses to submergence and flood stress. The epitranscriptomics response on flood stress wasn’t studied so far, but the opposite, drought stress, decreased the number of m6A modified sites in *Hippophae rhamnoides* (Zhang *et al*., 2021). Lowering methylation of drought stress response genes was also observed in poplar, but opposite effect was noticed in tobacco (Xiang *et al*., 2024). However, the patterns of epitranscriptome modification under water related stresses are still unexplored in plants, especially in the context of cytosine methylation and Ψ synthesis. Our study revealed that the proportion of m5C- modified cytosines to total cytosines was determined to be 0.08% in aquatic environment and 0.03% in terrestrial environment. Similar proportions were observed in a study using *A. thaliana*, where the highest value (0.028%) was found among different RNA types (Cui *et al*., 2017). In mRNA from siliques and rosette leaves, this coefficient was highest at 0.036% and lowest at 0.01%. The abundance of m5C was also found to increase after 15 days of cultivation, from 0.027% on day 3 to 0.033% in seedlings (Cui *et al*., 2017). Transcripts exhibiting significant m5C modifications, identified in the *R. fluitans* transcriptome from both aquatic and terrestrial environments, were associated with various responses to stimuli, including ’response to stimulus,’ ’response to stress,’ ’response to oxygen-containing compounds,’ and ’response to abiotic stimuli’. Previous research on various rice tissues (Yu *et al*., 2023) has been corroborated by our findings, which demonstrated that m5C-modified transcripts were involved in stress responses. Differences in the proportion of m6A- modified nucleotides to unmodified adenines were observed. In terrestrial *Riccia*, this proportion was found to be 0.04%, while in aquatic *Riccia*, it was determined to be 0.1%. In *A. thaliana*, the level of m6A relative to unmodified adenines varied across different tissues, reaching 0.9% in leaves, 0.6% in roots, and 1.5% in young seedlings (Zhong *et al*., 2008). Similar to the m5C modification, transcripts with a significant m6A modification in both environments were involved in responses to stimuli such as ’response to stimulus’, ’response to stress’, and ’response to oxygen-containing compound’. Differentially methylated peaks in *Arabidopsis* were involved in defense response pathways in response to temperature changes (Wang *et al*., 2023). In *R. fluitans*, the lowest proportion of Ψ modifications was found to be 0.001% in terrestrial forms and 0.002% in aquatic forms, although literature suggests considerable variability. Transcriptome-wide analysis revealed 323 transcripts containing Ψ, a number comparable to those reported in *A. thaliana* (Sun *et al*., 2019), despite employing different detection methods. Transcripts with the detected Ψ modification were involved in GO processes related to stress and photosynthesis, similar to the transcripts identified in *Arabidopsis* (Sun *et al*., 2019). The identification of Ψ sites in human mRNA was found to vary significantly across three different publications, ranging from hundreds to thousands, with only a small subset of these sites being common to all three studies (Carlile *et al*., 2014; Schwartz *et al*., 2014; Li *et al*., 2015). In conclusion, the detection of Ψ sites within the same species can yield highly variable results (Ramakrishnan *et al*., 2022). The most significant challenge is likely to be the accurate determination of Ψ modification sites. Till now it was also unknown whether the same mRNA molecule carries the Ψ, m6A, and m5C sites (Ramakrishnan *et al*., 2022), but in *R. fluitans* 304 transcripts with such feature was identified.

One of the aspects investigated in our study was the impact of RNA modifications on transcript length. The results indicated that transcripts containing the detected Ψ modification were surprisingly shorter compared to other isoforms of the same gene. An exception was observed for transcripts with the m5C modification in an aquatic environment, which exhibited longer products. Additionally, a shortening of poly(A) tails was observed in transcripts with detected RNA modifications under flood conditions and in transcripts with Ψ in terrestrial *Riccia*. Shortened poly(A) tails were also detected in rice transcripts with m5C and m6A modifications (Yu *et al*., 2023). It is important to note that the poly(A) tail is not a static feature of the transcript but rather a dynamically changing structure (Jalkanen *et al*., 2014). Interestingly, transcripts with detected Ψ in an aquatic environment showed a similar trend in poly(A) tail length and overall transcript length based on exon and intron content. It was observed that gene length might correlate with its expression frequency. Longer genes were found to be less frequently used after development, while shorter genes were often found to play a significant role in the organism’s daily functioning, such as in the human immune system (Lopes *et al*., 2021). Unfortunately, no extensive research has been conducted to correlate gene length with function in plants. Studies investigating the length of cDNA in *Arabidopsis* seedlings subjected to heat stress revealed a higher translation efficiency for longer molecules (Yángüez *et al*., 2013). The possibility was raised that ribosomes remained bound to long mRNAs for a longer period and that translation patterns could be modulated by organisms through the control of ribosome density under stress (Ingolia *et al*., 2009; Piques *et al*., 2009; Wu *et al*., 2012). Studies have shown that the proximity of the 3’ end to the ribosomal recruitment site of mRNA could lead to preferential translation of shorter transcripts (Fernandes *et al*., 2017). In our study, alternative splicing in modified transcripts was investigated. This process was hypothesized to be correlated with transcript length. An increased retention intron (RI) was demonstrated in transcripts with detected modifications compared to other isoforms of the same genes. It has been shown that m6A modification both promoted and inhibited alternative splicing events in sex determination, reproductive development, inflammatory response, and adipogenesis in mammals (Kasowitz *et al*., 2018; Wang *et al*., 2018; Huang *et al*., 2021*a*; Li *et al*., 2022). m6A was found in alternative introns/exons of mouse embryonic stem cells, suggesting a potential correlation between m6A and RNA splicing (Wei *et al*., 2021). Ψ in pre-mRNA has been shown to regulate splicing by influencing the secondary structure of pre-mRNA (Sousa-Luís and Carmo-Fonseca, 2022). Unfortunately, there is a lack of publications regarding the direct impact of m5C on alternative splicing. Undoubtedly, a correlation exists between RNA modifications (Ψ, m5C, m6A) and alternative splicing, which may prove to be an interesting topic for future scientific research.

The identification of a core set of de-/methyltransferases and Ψ synthases conserved between *R. fluitans* and the model plant *A. thaliana* provides valuable insights into the evolution of RNA modification machinery and underscores the fundamental importance of these enzymes in regulating gene expression across land plants. The presence of *RfMETTL3* and *RfFIP37*, key components of the m6A methyltransferase complex (Bodi *et al*., 2012; Garcias Morales and Reyes, 2021), highlights the conserved role of m6A modification in *R. fluitans*. The identification of *RfALKBH, RfALKBH9B*, and *RfTET8* further emphasizes the dynamic nature of RNA modifications, with these enzymes likely contributing to active RNA demethylation in *R. fluitans*. The absence of *RfALKBH, RfALKBH9B*, and *RfNSUN2* homologs in the moss *P. patens* raises questions about gene loss or divergence in the moss lineage. The epitranscriptomics of model moss is poorly explored, but m6A methylation seems to play an important role in phenotypic plasticity of *P. patens* by contribution to the transition from 2D to 3D growth (Garcias-Morales *et al*., 2023). All main three m6A writers are present in *P. patens*, however we weren’t able to identify m6A erasers coding genes, which require further investigation. The absent NSUN2 family proteins are identified as m5C writers (Zhang *et al*., 2023*b*) among two others writers, DNMT2 and TRDMT1, which are present in *P. patens*. This observation could reflect differences in the regulatory networks or functional specialization between liverworts and mosses. Opposing the m5C ‘writer’ proteins (DNMT2, TRDMT1, NSUN2) are the ten–eleven translocation (TET) enzymes, functioning as ‘erasers’, which have demonstrated their ability to catalyze the demethylation of m5C (Fu *et al*., 2014; Delatte *et al*., 2016). In this study only one TET homolog was identified in liverworts, in comparison to three in *P. patens* and 15 in *A. thaliana*. Angiosperms have undergone extensive gene duplication events throughout their evolutionary history, which allows for duplicated genes to evolve new functions or specialize in existing ones (Crow and Wagner, 2006; Panchy *et al*., 2016). TET genes may have duplicated and diversified in angiosperms, leading to the larger number of homologs which are involved also in DNA demethylation processes (Ji *et al*., 2018). However, it is important to note that both *R. fluitans* and *M. polymorpha* belong to the complex thalloid liverworts, and it remains unknown whether other liverwort lineages possess a greater diversity of TET genes. This is particularly relevant considering that most complex thalloid liverworts lack some RNA modification mechanisms, such as RNA editing, which are present in other liverwort lineages (Rüdinger *et al*., 2008; Dong *et al*., 2019; Myszczyński *et al*., 2019). Therefore, while the reduced number of TET genes in *R. fluitans* may suggest lineage-specific adaptations and potential functional divergence (Fig. 6), further investigation across a broader range of liverwort diversity is needed to fully understand the evolutionary history of these enzymes in land plants. The orthologues number of Ψ writers are more conserved, especially among bryophytes (Fig. 7) with presence of all six PUS families (Spenkuch *et al*., 2015). Our results suggest that while a core set of enzymes is conserved across land plants, lineage-specific adaptations have led to variations in gene family size and potentially functional divergence. Further research is needed to functionally characterize the identified de-/methyltransferases and Ψ synthases in liverworts and to determine their specific RNA targets. Comparative transcriptomic and epitranscriptomics studies across different plant lineages could provide further insights into the correlation between RNA modification patterns and gene expression profiles. Ultimately, a deeper understanding of RNA modification pathways in early-diverging land plants like *R. fluitans* will shed light on the evolution of gene regulation mechanisms in plants and their adaptation to diverse environments.

Despite differences in transcriptome-wide methylation and pseudouridylation between the two environments, the expressions of genes coding enzymes involved in studied modifications are not altered (Table S35). This observation raises questions about the direct correlation between the expression of these enzymes and the global levels of RNA modifications. The findings challenge the notion that global mRNA modification levels can be reliably predicted solely based on the expression of writer/eraser enzymes (Zhong *et al*., 2008; Shen *et al*., 2016; Cui *et al*., 2017) and Ψ synthetases (Xie *et al*., 2022). While enzyme expression provides valuable insights into the potential for modification, it does not necessarily reflect the actual modification status of RNA transcripts. The discrepancy between enzyme expression and modification levels underscores the need for direct measurement of RNA modifications using techniques like nanopore direct RNA sequencing (Yu *et al*., 2023; Maździarz *et al*., 2024). However, the identification and quantification of m6A via DRS, particularly in the context of diverse sequence motifs, remains a subject of ongoing investigation and methodological refinement. Traditionally, m6A detection has been focused on the DRACH motif (where D = A, G, or U; R = A or G; and H = A, C, or U), a sequence context recognized by the methyltransferase complex responsible for installing m6A (Hendra *et al*., 2022; Huang *et al*., 2024). The recent studies on plant epitranscriptomes suggested that m6A modifications can also occur in non-DRACH contexts, expanding the potential scope and complexity of m6A-mediated gene regulation (Hou *et al*., 2022). In plants, m6A sites are primarily found near the stop codon and within the 3’ UTR, and they predominantly display the m6A consensus motif RRACH, along with other motifs like URUAY (Wang *et al*., 2023). Nanopore direct RNA-seq employed to map m6A sites at a single-base resolution in *Arabidopsis* revealed that the DRAYH motif was enriched at m6A sites, and AAACU and AAACA were the most commonly observed motifs (Parker *et al*., 2020). Recent advances in analyzing raw nanopore signals and basecalling models enabled transcriptome-wide, all-context (AC) detection of m6A (Acera Mateos *et al*., 2024). This approach revealed a substantially higher number of m6A sites compared to DRACH-only analyses, highlighting the potential underestimation of m6A methylation levels when focusing solely on the DRACH motif (Maździarz *et al*., 2024). The discrepancy between DRACH-only and AC analysis underscores the importance of considering the broader sequence context in which m6A modifications can occur. The findings of this study, along with observations in other organisms, emphasize the importance of adopting a comprehensive approach to m6A detection and quantification. By considering both DRACH and non-DRACH motifs, researchers can gain a more accurate and complete understanding of the m6A epitranscriptome and its role in gene regulation. This knowledge is particularly crucial for studying non-model organisms like *R. fluitans*, where the epitranscriptome landscape is less well-characterized.

Detection of RNA bases modification using DRS is in rapid development, especially in the case of adenine methylation (Hendra *et al*., 2022; Acera Mateos *et al*., 2024; Huang *et al*., 2024). However, the Ψ and m5C identification models for R9.4.1 pore are limited (Huang *et al*., 2021*b*), but can be validated using TGS with adapted PRAISE method which allow for the identification of Ψ positions through the appearance of deletions at the site of this modification as well as m5C via C/T conversion ratio. Deletions at Ψ positions in *R. fluitans* transcripts from an aquatic environment were identified using the PRAISE method, thereby confirming the presence of Ψ at these sites. Applied method also enabled validation of m5C sites identified by DRS. These results support the application of third-generation sequencing in conjunction with the PRAISE method for accurate mapping of pseudouridylation and m5C sites (Zhang *et al*., 2023*a*) using high-throughput and more accurate R10.4.1 flow cells. Recent improvements of nanopore dorado basecaller with connection of latest RNA004 chemistry enabled detection of wide range RNA modifications like m6A, m5C, inosine and Ψ bases with high, all context precision and over 99% confidence [https://github.com/nanoporetech/dorado].

## Acknowledgements

The study was financially supported by The National Science Center Kraków, Poland: Grant No. 2020/39/B/NZ8/02504.

ORCID: Mateusz Maździarz (ORCID 0000-0002-7278-4604) Katarzyna Krawczyk (ORCID 0000-0001-9019-0490) Joanna Szablińska-Piernik (ORCID 0000-0003-0265-940X) Łukasz Paukszto (ORCID 0000-0002-3618-1064) Monika Szczecińska (ORCID 0000-0002-5377-4304) Paweł Sulima (ORCID 0000-0002-8290-8180) Jakub Sawicki (ORICD 0000-0002-4759-8113) Conceptualization – Mateusz Maździarz, Jakub Sawicki Investigation – Katarzyna Krawczyk, Joanna Szablińska-Piernik, Paweł Sulima Software - Mateusz Maździarz, Łukasz Paukszto, Jakub Sawicki Formal analysis - Mateusz Maździarz, Łukasz Paukszto, Jakub Sawicki Methodology – Mateusz Maździarz, Łukasz Paukszto, Katarzyna Krawczyk, Joanna Szablińska- Piernik, Paweł Sulima, Jakub Sawicki Supervision – Jakub Sawicki Writing – original draft – Mateusz Maździarz, Łukasz Paukszto, Katarzyna Krwaczyk, Joanna Szablińska-Piernik, Paweł Sulima, Monika Szczecińska, Jakub Sawicki Writing – review & editing – Mateusz Maździarz, Jakub Sawicki Project administration – Jakub Sawicki Funding acquisition – Jakub Sawicki

---

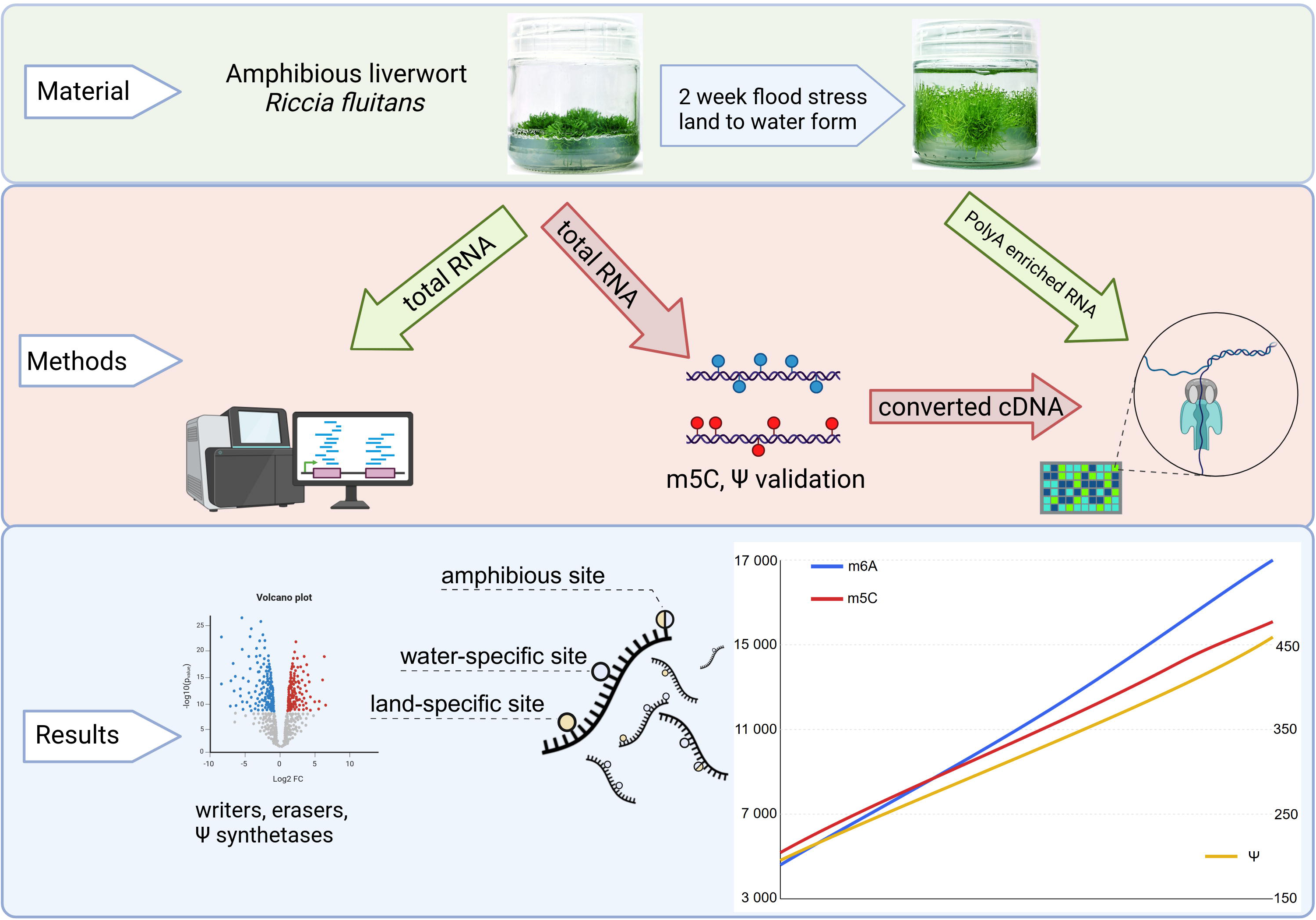

